# Resource-use plasticity governs the causal relationship between traits and community structure in model microbial communities

**DOI:** 10.1101/2024.08.06.606555

**Authors:** Brendon McGuinness, Stephanie C. Weber, Frédéric Guichard

## Abstract

Resolving the relationship between species’ traits and their relative abundance is a central challenge in ecology. Current hypotheses assume relative abundances either result from or are independent of traits. However, despite some success, these hypotheses do not integrate the reciprocal and feedback interactions between traits and abundances to predictions of community structure such as relative abundance distributions. Here we study how plasticity in resource-use traits govern the causal relationship between traits and relative abundances. We adopt a consumer-resource model that incorporates resource-use plasticity that operates to optimize organism growth, underpinned by investment constraints in physiological machinery for acquisition of resources. We demonstrate that the rate of plasticity controls the coupling strength between trait and abundance dynamics, predicting species’ relative abundance variation. We first show how plasticity in a single species in a community allows all other non-plastic species to coexist, a case of facilitation emerging from competitive interactions where a plastic species minimizes its similarity with competitors and maximizes resource-use efficiency in its environment. We apply this environment-competition trade-off to predict trait-abundance relationships and reveal that initial traits are better predictors of equilibrium abundances than final trait values. This result highlights the importance of transient dynamics that drive species sorting. The temporal scale of transients determines the strength of species sorting due to the emergence of ‘ecological equivalence’ at equilibrium. We propose trait-abundance feedback as an eco-evolutionary mechanism linking community structure and assembly, highlighting trait plasticity’s role in community dynamics.

## Introduction

Understanding the relationship between species traits and community structure, the variation in relative abundances among species, remains an important challenge in ecology. The connection between species traits and their relative abundances can elucidate the ecological and evolutionary processes responsible for the maintenance of species and functional diversity. Traditional niche-based theories of community assembly assume a strong trait-abundance relationship underpinned by the relationship between traits and habitat characteristics (1–3), while neutral theory assumes no relationship between traits and abundances (4). Both of these approaches ignore the well documented feedback between traits and abundances, which can occur either genetically via eco-evolutionary processes (5, 6) or through phenotypic plasticity (7). However, we still lack a general theory integrating these feedbacks into predictions of community structure and its relationship with species traits. Here we study resource-use plasticity in model microbial communities to determine the role of plasticity rate for the emergence of community structure and its relationship with species traits.

In contrast to the niche-based view which links traits and structure through species sorting, neutral theory provides an explanation for species relative abundances through stochastic processes such as drift and dispersal (4). In this framework, it is assumed that species are ‘ecologically equivalent’, meaning their traits have no correlation with their abundances (4). Rather than being mutually exclusive, many have posited that niche and neutral frameworks define the extremes of a continuum along which ecological reality exists (8–11). However, within this continuum framework, we are caught in a paradigm that explains community assembly from the extent to which traits affect structure, ignoring the feedback from structure onto traits. By incorporating explicit feed-backs between traits and abundance, plasticity could help explain trait-abundance relationships and ecological equivalence as emerging properties of coupled dynamics between traits and abundance rather than resulting from the balance between alternative mechanisms of species interactions.

Ecological and evolutionary processes have been shown to act on species that are not initially ecologically equivalent, but dynamically converge on states that are, a phenomenon referred to as emergent neutrality, where abundances disassociate from traits (12–14). Recent studies on model microbial communities show ecological equivalence emerging in niche-based models when constraints are imposed on the physiological machinery used for the acquisition of resources (15– 17). Subsequent work demonstrates that resource-use plasticity promotes diversity when species are allowed to change their resource uptake rates to maximize their own growth (18). Here, species self-organize to always reach an equilibrium in which species become ecologically equivalent, where ‘emergent neutrality’ does not depend on fixed resource uptake rates of species in a community. Although these studies focus on resource-use trait plasticity in microbes, plasticity in resource-use is ubiquitous and has for example been investigated under the umbrella of adaptive foraging (19) in systems such as plant-pollinator networks (20), arthropods (21), and plants (7, 22). The positive effect of trait plasticity on species diversity has been shown in theoretical, experimental and field studies (18, 23, 24). However, it remains unclear whether plasticity, or more generally trait-abundance feed-backs, govern the emergence of community structure and its causal relationship with species traits.

Here, we propose plasticity as a mechanism that governs the trait-structure relationship by controlling the transient time before ecological equivalence. By adopting a consumer-resource model that incorporates trait plasticity in resource-use, we reveal a feedback between the dynamics of abundance and of traits and the conditions that give rise to a predictable relationship between species traits and their relative abundances. We first isolate this trait-abundance feedback onto a single plastic species showing that plasticity in a single species can rescue a community and allow for the coexistence of multiple competitive species by minimizing both its similarity with competitors and its distance from the environment optimum. We then study communities where all species are plastic. We observe that initial traits predict equilibrium relative abundances better than equilibrium traits, and study how transient dynamics that result from initial abundances and traits drive species sorting. We further show that the temporal scale (rate) of plasticity controls not only the strength of the correlation between traits and abundances, but also the emergence of community structure itself. Our results reveal how the temporal scale of plasticity determines the strength of transient niche processes in model microbial communities. These findings offer an eco-evolutionary mechanism that links the processes that govern community structure and assembly, underscoring the importance of trait plasticity in ecological dynamics.

## Methods

### Niche-based model with metabolic trade-offs

We first overview a niche-based model of microbial communities in which ecological equivalence can emerge as the equilibrium state (15). In this consumer-resource model, it is assumed that all, *N*_*S*_, species (consumers) have the same total protein budget, *E*, for the uptake and metabolism of *N*_*R*_ substitutable resources. These resources are externally supplied at rates 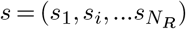 for *i* = 1, …, *N*_*R*_. Species maintain different fixed species-level allocations of these proteins that confer differences in rates of resource-use, to which we will refer as species’s traits. Each species *σ* uptakes resource *i* at a rate *α*_*σi*_ ≥ 0. We assume that resource uptake rate, *α*_*σi*_, is proportional to species *σ*’s proteomic investment in the uptake and metabolism of resource *i*. Following (15), this metabolic trade-off is imposed with the constraint 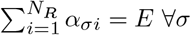, leading to:

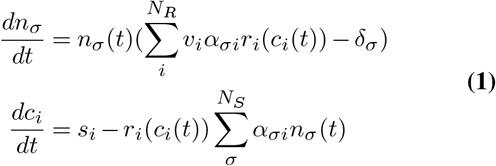

where *n* is the abundance of species *σ*, and *c* is the concentration of resource *i*. Here we take the mortality or maintenance rate to be the same for all species such that *δ*_*σ*_ = *δ* ∀*σ*. The nutritional value *v*_*i*_ is also taken to be the same over all resources such that *v*_*i*_ = *v* ∀*i*. This model uses a Monod function to describe resource-use as 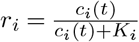 where *K*_*i*_ is the half saturation constant for resource *i*.

In the absence of metabolic trade-offs, the diversity of the system has an upper bound of *N*_*R*_ set by competitive exclusion. However, the metabolic trade-off implies that when all resources (scaled by nutritional values if 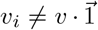) are available in equal concentrations, there is a point where the zero net growth isoclines (ZNGI) for all consumers intersect, meaning that all consumers will have equal fitness (taken as growth rate) and coexistence of all *N*_*S*_ species is possible (25). The equal resource concentration requirement is met when the normalized supply vector 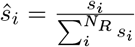 is in the convex hull of the normalized resource-uptake traits, 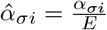, of all *σ* species in the community (15). In this scenario, resource concentrations are driven to equal values (assuming equal nutritional value *v*_*i*_) at equilibrium, and all species have the same fitness, resulting in the emergence of ecological equivalence among all *N*_*S*_ species. Importantly, when the aforementioned convex hull condition is met, the abundances equilibrium is not limited to a single fixed point but is instead possible on an infinite set of fixed points (mathematically degenerate) set by initial conditions, on a neutral manifold. We refer to the normalized supply vector as the supply vector for short in the rest of this manuscript.

### Niche-based model with resource-use plasticity

Microbes are known to change their resource-use based on the quantity and quality of resources in the environment (26), and via gene regulation have the ability to make these choices depending on what resources can maximize growth rate (27, 28). Recent work has incorporated resource-use plasticity to the consumer-resource model (Eq. 1) (18). In this case, the resource uptake rates, *α*_*σi*_, are dynamic and considered as state variables in the system of differential equations (Fig 1a). Resource uptake rates 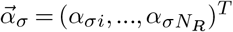 change in time to maximize the fitness of species *σ* which is measured as the growth rate 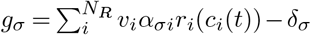. This growth rate optimization is implemented by gradient ascent in which the consumption strategies follow the gradient ascent equation:

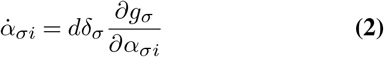

where *d* is the plasticity rate setting the velocity of the dynamic resource-use adaption. In this equation, the resource uptake rates are unbounded, meaning that they will just increase towards infinity if no constraint is added. We know however, that microbes have a limited capacity towards their enzymes and uptake proteins used towards resource uptake rates (29). To incorporate this biological realism, a constraint in the form of an upper bound which imposes a maximum total resource uptake rate 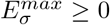 for each species *σ* is included in this model. Here we assume that all species are on a ‘Pareto frontier’, meaning that 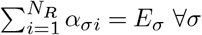. So long as the ratio between total resource uptake and maintenance rates is the same across all species (see (18) and SI for justification), 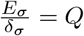 ∀*σ*, then all *N*_*S*_ species can coexist independently from the convex hull condition on initial traits from the non-plastic trait model (equation (1)). Instead, the community ‘self-organizes’ to an equilibrium state in which the convex hull condition is met. However, if plasticity rate, *d*, is too slow then exclusion occurs on a timescale faster than species traits can acclimate to a coexistence state. Now an infinite set of fixed points exist for an infinite possible set of equilibria *α*_*σi*_ all depending on initial values of *α*_*σi*_ and *n*_*σ*_.

**Fig. 1.**
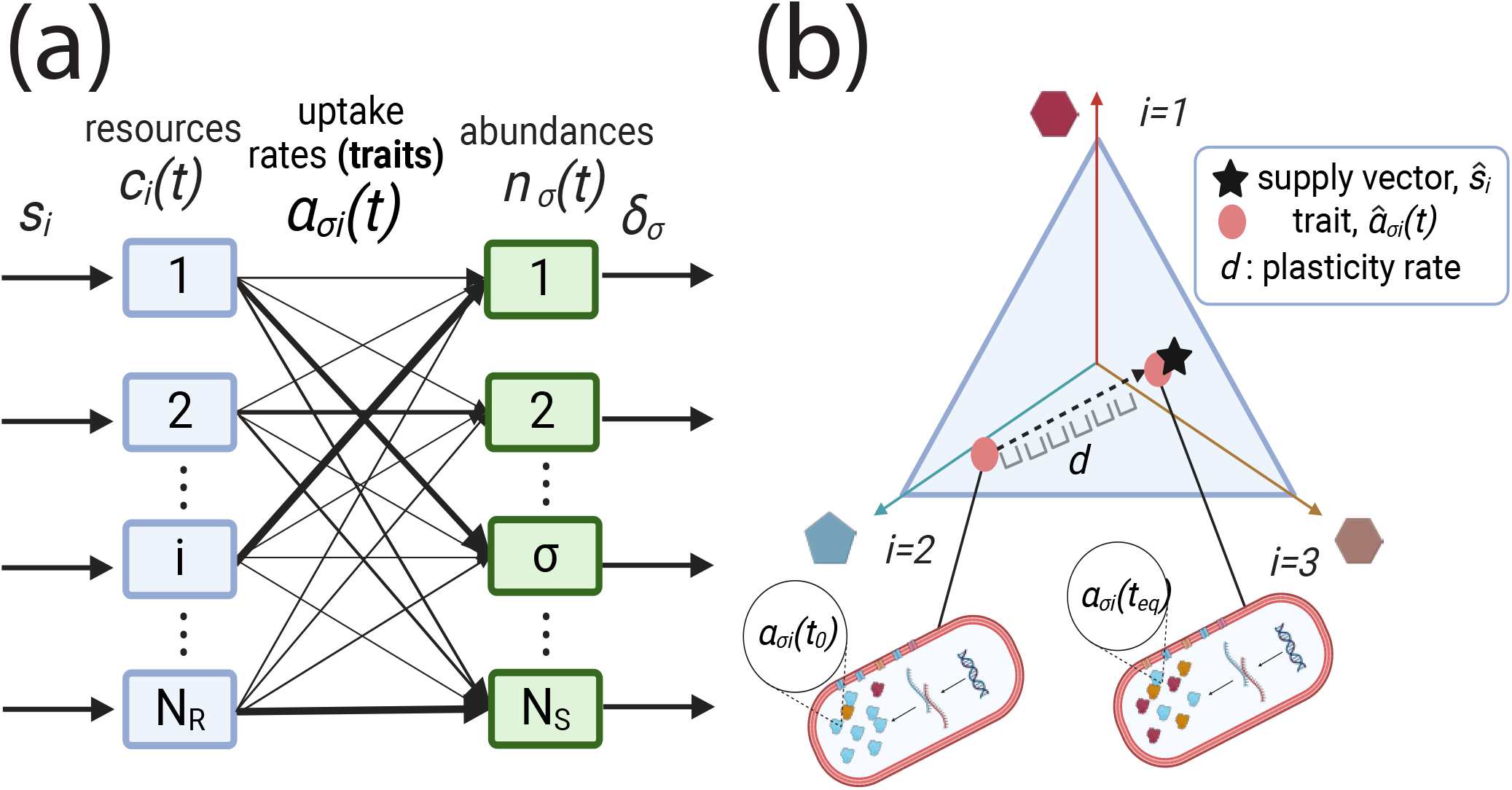
Model schematic. (a) Schematic of consumer-resource model where resource-use is plastic. Weights of uptake edges (traits) are dynamic and based on growth optimization by gradient ascent in equation (2). (b) Simplex diagram where a species’ traits (normalized resource-use strategy, 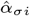) change based on *N*_*R*_ = 3 resources available in the environment. The supply vector (black star, *ŝ* _*i*_) is normalized and plotted on the same simplex.

### Inferring relative abundances from traits

In order to move past the binary description of coexistence or exclusion, we build a statistical model to explain variation in equilibrium community structure, 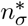 (* denotes equilibrium), from species traits, *α*_*σi*_(*t*) at a given time *t*. Our model is derived from the fact that species are trying to maximize their growth with plastic resource-use traits (uptake rates), as assumed in the model we adopt. In the plastic consumer-resource model, species maximize their growth in two ways: (1) maximize their distance in trait space on the simplex from competitors and (2) minimize their distance to the supply vector. However, in order to calculate distances, we first must convert our *N*_*R*_ − 1 dimensional-simplex (i.e. barycentric coordinate system) into Cartesian space. This transformation is described by the *N*_*R*_*×N*_*R*_ − 1 dimensional matrix

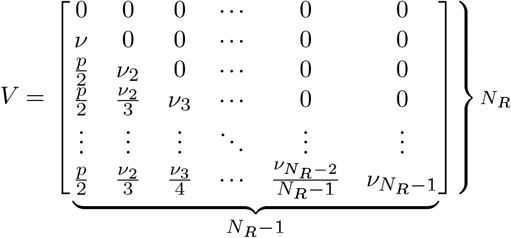

where *ω* is the side length of the simplex, which we take to be 1, and

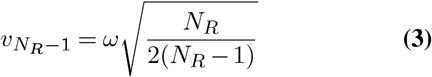

is the height of an *N*_*R*_ − 1 dimensional simplex. By adding this height to the centroid of an *N*_*R*_ − 2 dimension simplex, we can recursively construct the *N*_*R*_ − 1 dimensional simplex where the rows of *V* denote the Cartesian coordinates for vertex *i*. With the coordinates of these vertices, we can then convert any barycentric point, such as our normalized resource uptake rates, 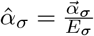, and our normalized supply vector, 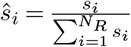, into a Cartesian point. For resource uptake rates, this occurs through the transformation 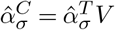 where *C* denotes that vector is in Cartesian space and *T* is the transpose of the vector in barycentric space (i.e. simplex). Likewise, for the normalized supply vector the, transformation is 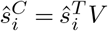. We note that the simpler way of describing a simplex in Cartesian space is by transforming the point with the identity matrix 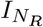, such that 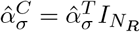. However, this represents the simplex in an extra dimension which prevents us from being able to numerically calculate the convex hull using the `ConvexHull` function in the package `scipy.spatial`.

With these Cartesian points of relative traits and resources, we can calculate distances between traits using the *L*^2^-norm. We define a *N*_*S*_*×N*_*S*_-dimension pairwise trait distance matrix as 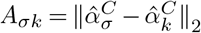 for species *σ, k* = 1, …, *N*_*S*_ where *C* denotes Cartesian space and and ∼ denotes relative trait, abundance or resource levels. Additionally, we define a vector of length *N*_*S*_ of the distances between the traits of species *σ* and the normalized supply vector *ŝ*_*i*_ described as 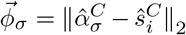. Since growth maximization attempts to maximize trait distance from competitors and minimize trait distance to supply vector, we use these two distances as inputs into a statistical model that predicts the abundance of species *σ* as

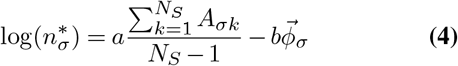

where parameters *a* and *b* are fitted by ordinary least squares regression. We use *R*^2^ as the performance metric of our statistical niche-based model in predicting equilibrium abundances. The first term is scaled by *N*_*S*_ − 1 to preserve the relative scales of parameters *a* and *b* which correspond to strength of the terms associated with avoiding competitors (biotic) and and maximizing resource efficiency (abiotic). We report on these relative strengths as a function of plasticity rate in the SI. We compare this niche-based model to the null expectation assuming no species sorting, 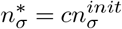 where every species grows in equal proportion to their starting abundances such that there are no fitness differences generated by traits.

### Quantification of community structure

We calculate the trait-driven variation in equilibrium abundance among species as the complement of the normalized Shannon Entropy, which we refer to as the Species Sorting Index (SSI) described as:

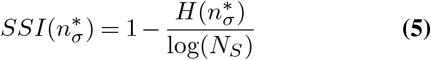

where 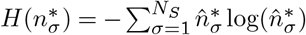 describes the Shannon Entropy with 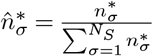 being the relative species abundance distribution at equilibrium. We assume species’ abundances are initially equal, such that 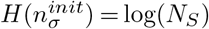 and 0 ≤ *SSI* ≤ 1. With this assumption, we are able to isolate the variation in relative abundances driven by heterogeneous traits and subsequent trait-abundance feedback interactions. Here, *SSI* = 0 if there is no trait-driven variation of abundances, and *SSI* = 1 for maximum trait-driven variation of abundances, i.e. if a single species excludes all the others.

### Calculation of community distance from supply vector

The strength of species sorting depends on how far a community begins from the system’s attractor, the supply vector *ŝ*_*i*_. We calculate this quantity as the centroid-to-supply vector distance, which we define as

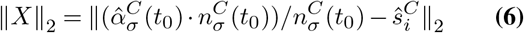

where *t*_0_ = 0. This quantity is calculated as the *L*^2^-norm between the trait centroid weighted by abundance, which can be thought of as the center of mass of the traits if each point is weighted by its respective abundance, and the supply vector. To calculate both the weighted-centroid and its distance from the supply vector, these quantities are converted to their Cartesian coordinates in the method described in the previous section.

## Results

### Community rescue from a single plastic species

To better understand the coupling between traits and abundances, we first investigate the scenario in which only one species is allowed resource-use plasticity while all the other species have fixed traits. This approach allows us to isolate the feedback from species abundances onto the dynamic traits of a single species. Previous work shows coexistence of *N*_*S*_ *> N*_*R*_ is possible if the normalized supply vector (which we refer to as supply vector for succinctness), *ŝ*_*i*_, is within the convex hull of all the traits in a community,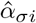 (15). We observe that one plastic species in a community is sufficient for reaching coexistence. To illustrate this, we show a *N*_*S*_ = 10 species community competing for *N*_*R*_ = 3 resources on a simplex plot where the supply vector (denoted by the black star) is outside of the initial convex hull (area enclosed by blue line) of the competing species’ resource traits (Fig 2a). If the convex hull condition is not met with fixed traits (blue line), competitive exclusion occurs (Fig 2b). However, when allowing one species (denoted by the blue ‘x’ in Fig 2a) to plastically change their resource uptake rates to maximize their growth, all species have positive abundances at equilibrium, i.e. coexistence occurs (Fig 2c). Although the plastic species changes its resource uptake rates to maximize its own growth, these traits change due to two operating forces: repulsion from competitors and attraction to the supply vector. The former is due to a species minimizing similarity with competitors and the latter due to a species maximizing resource-use efficiency. In the absence of competitors, the best resource-use strategy is at the supply vector. The consequence of this trait-abundance feedback is that the static species ’push’ the plastic species away from the supply vector, making the final convex hull of the community always include the supply vector (area enclosed by orange line in Fig 2a). This plastic species thus rescues all of the species that would have been excluded.

**Fig. 2.**
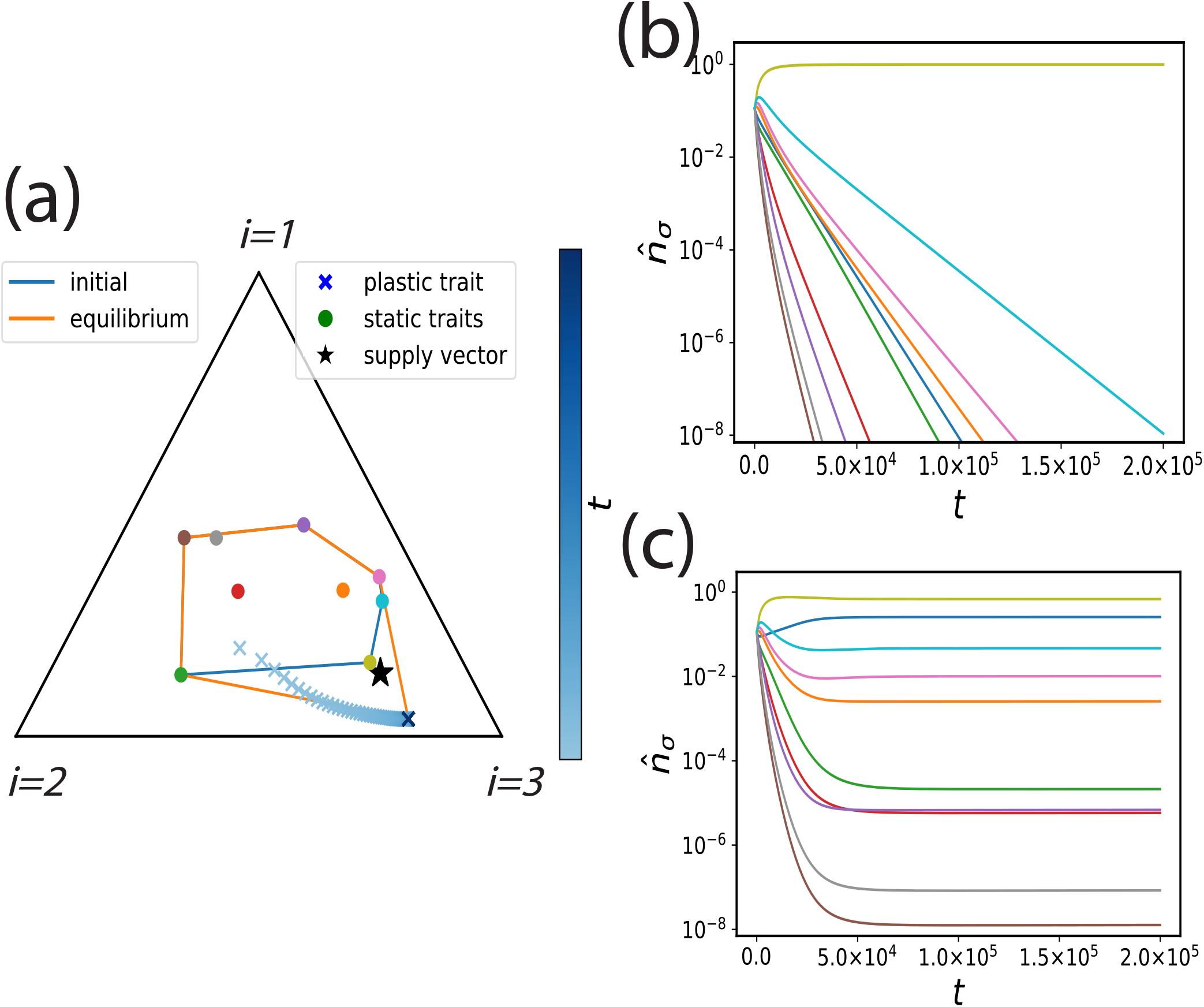
Plastic species promotes coexistence in community with *N*_*S*_ = 10, *N*_*R*_ = 3. (a) Trait time-series on simplex plot where plastic species’ traits through time are denoted by the ‘x’ and static species are denoted in circles. The blue line is the convex hull of the initial traits (or if all species are static) while the orange line is the convex hull of the equilibrium traits if one species is plastic. (b) Time-series when all uptake strategies are static denoted by blue convex hull of the initial traits (c) Time-series when one uptake strategy is plastic, denoted by the blue ‘x’ and orange convex hull in (a). See SI for rest of parameters.

An important caveat of this plasticity-induced coexistence is that the species in the community have relative abundances that span eight orders of magnitude (Fig 2c). Although species are technically coexisting, the distribution of species abundances in this community is quite uneven. Even if the convex hull eventually finds its way to envelop the supply vector, some species could eventually fall under some extinction threshold. Thus, we focus our subsequent analysis on exploring what processes explain the variation in relative abundances as a direct result from traits-abundance feedbacks in plastic communities.

### Initial traits explain equilibrium relative abundances better than equilibrium traits

We extend our investigation to communities where all species exhibit plastic resource-use strategies to elucidate the contribution of variation in species traits to variations in relative abundances. Our single plastic species scenario in Fig. 2 demonstrates that trait plasticity operates to maximize growth in two ways: (1) maximize distance in trait space on the simplex from other competitors and (2) minimize their distance to the supply vector. We now predict equilibrium species abundances from these trait distances using either initial or equilibrium traits as inputs (Fig 3a,b). We build our predictions from a statistical model that assumes the log-transformed equilibrium abundance of each species is a linear combination of its trait distance from competitors (Fig 3a,b in dotted black lines) and of its distance to the supply vector (Fig 3a,b in dashed red line). We used this statistical model (equation (4)) to predict equilibrium relative abundance from initial or equilibrium trait values. We observe that initial traits are better predictors of equilibrium relative abundance than equilibrium traits (Fig 3c). Species traits lose their predictive capacity of equilibrium relative abundance through time, suggesting that variation in relative abundances at equilibrium results from the transient trait and abundance dynamics. These changes in traits through time are a direct result of the feedback between initial traits and abundances. The important consequence of this result is that it implies that species trait history (initial conditions) plays a role in determining equilibrium relative abundances.

**Fig. 3.**
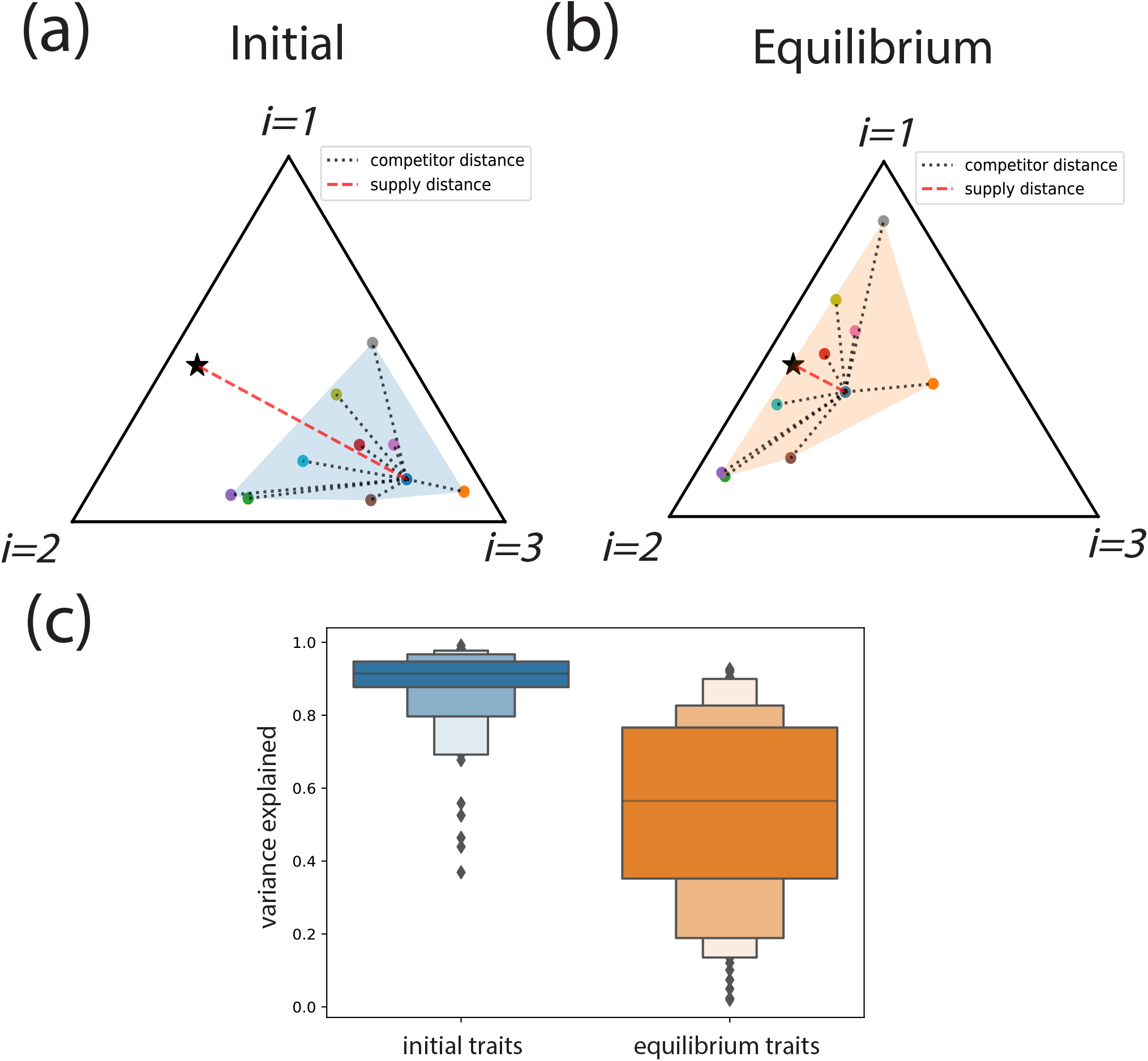
Initial traits are better than equilibrium traits at explaining variance in equilibrium abundances (a) Simplex of initial traits (blue) of a community of *N*_*S*_ = 10 and *N*_*R*_ = 3 illustrating the two terms of statistical model in equation (4) with competitor distance *A*_*σk*_ (black dotted), and supply distance *ϕ*_*σ*_ (red dashed) for species *σ, k* = 1,…, *N*_*S*_. (b) simplex of equilibrium traits (orange) illustrating the two terms of our statistical model with equilibrium traits as predictors. (c) Variance in equilibrium relative abundances explained (shown as *R*^2^) from equation (4) predicting equilibrium abundances from initial (blue) and equilibrium (orange) traits. See SI note 3 for rest of parameters.

These results suggest that near-equilibrium traits do not explain much variation of relative abundances. To assess the role of traits as drivers of relative abundances distribution near equilibrium, we perturb a community at equilibrium by randomly shuffling the abundances on to different traits and re-running the simulation (Fig. SI 1a, b). This perturbation pushes the trait centroid weighted by abundances off of the supply vector which serves as the community-level attractor (where the community converges to at equilibrium). When running a subsequent simulation that is ‘near equilibrium’, initial perturbed abundances predicted equilibrium abundances far better than traits (Fig. SI 2c), demonstrating weak sorting close to the equilibrium, in which near ’ecological equivalence’ emerges. With the predictability of species equilibrium relative abundances depending on how far the initial traits are from the community level attractor, we posit that the transient time of our system, largely set by the plasticity rate of the traits, will determine how well initial traits predict equilibrium relative abundances, i.e. how strong sorting acts on species.

### Plasticity rate drives long term impacts of trait-abundance transient dynamics

We now investigate the role of transient dynamics in our system by tracking transient times (defined as the time before equilibrium is reached) of traits and abundances over different plasticity rates, *d*. We observe that both transient time of species traits and abundances decrease with increasing *d*. However, as *d* is further increased, trait transient time continues to decrease while abundance transient time plateaus (Fig 4a). Because plasticity is the process bringing the community centroid to ecological equivalence over the supply vector, increasing *d* will limit the time spent under transient trait-driven ecological selection. In other words, this timescale, largely set by *d*, governs the strength of species sorting. We test this prediction with our statistical model to estimate the variation in equilibrium relative abundance that we can explain from initial traits (equation (4)) with increasing plasticity rate. We ran 300 simulations with initial trait values, initial species abundances, and resource supplies drawn from random uniform distributions spanning three orders of magnitude (see SI for full parameter details). We obtain the variation in equilibrium relative abundances explained by initial traits, which we compare to a null model where variance in equilibrium abundances is not generated by traits, but explained solely by initial abundances (Fig 4b). In the null model, ecological equivalence is assumed throughout the entire time-series. For slow plasticity rates *d*, equilibrium abundances are better explained by traits while at fast *d* equilibrium abundances are better explained by initial abundances (Fig 4b). Transient time of trait and abundance dynamics also diverge over fast plasticity rates where the null initial abundance model outperforms the initial trait model (Fig 4a, b). These results indicate that traits predict abundances when transients are long due to slow plasticity, and lose their predictive power at fast plasticity resulting in short trait transient times and with longer abundance transients occurring under weak sorting, i.e. while traits have reached their fitness invariant equilibrium.

**Fig. 4.**
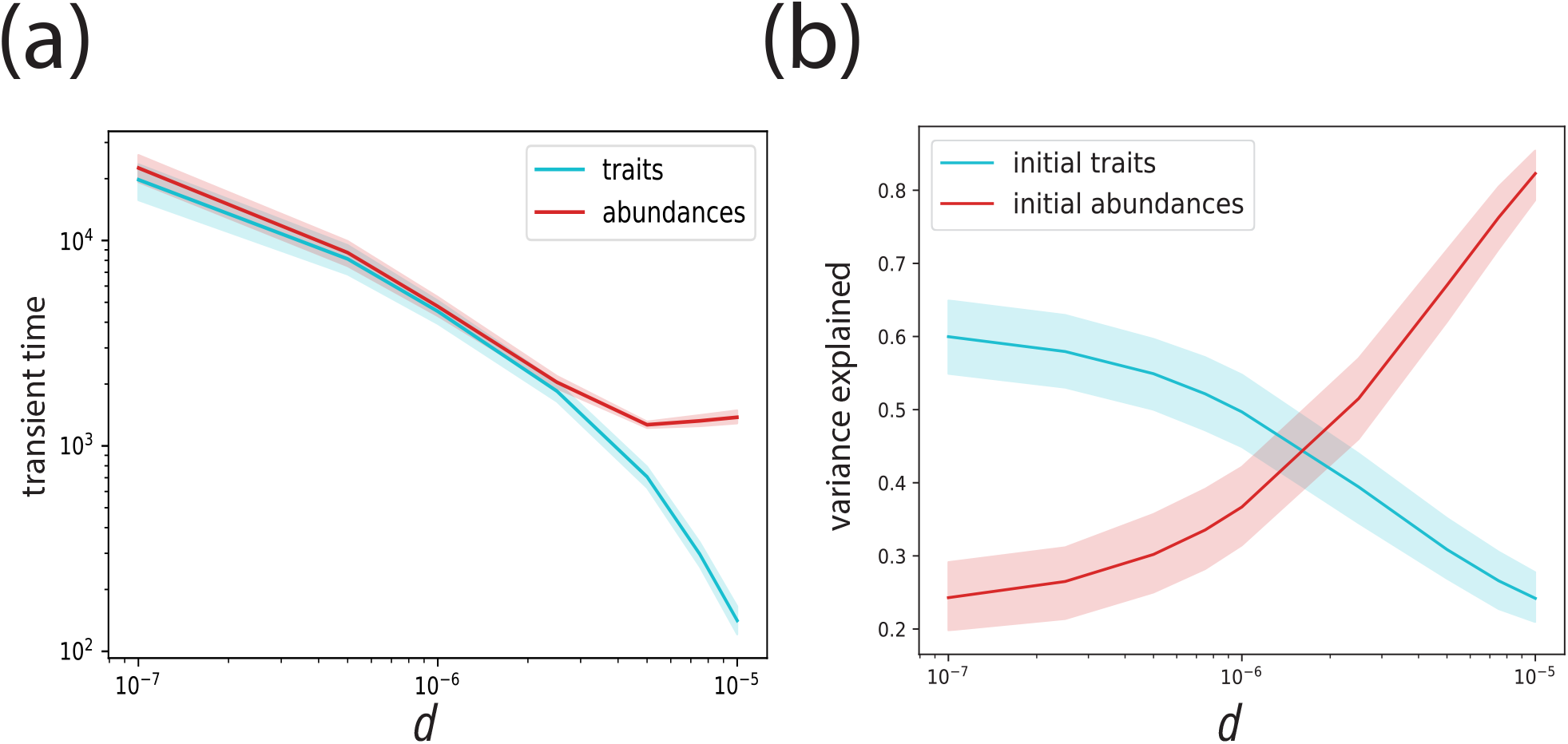
Plasticity rate *d* governs transient time in which trait-driven variation in community structure is generated. (a) Average transient time of traits (cyan) and abundances (red) in relation to *d* plotted with 95% confidence intervals. (b) Variance in equilibrium relative abundance explained (*R*^2^) by equation (4) with initial traits as input (cyan), and null initial abundance model (red), plotted with 95% confidence intervals. Examples shown for *N*_*S*_ = 10, *N*_*R*_ = 3. Initial relative trait, abundance, supply rates drawn from uniform distributions ranging 3 orders of magnitude. See SI for parameter details.

### Community structure emerges from transient fitness inequalities

So far our results indicate that transients control the strength of species sorting, i.e. how well species’ traits determine their relative abundance, in plastic competitive communities. Our previous results demonstrate that weak sorting occurs when the community centroid (the traits on the simplex weighted by their relative abundances) is near the supply vector, which acts as the community-level attractor of the system. These results suggest that sorting can remain strong during large enough transient times which are controlled by plasticity rate, *d*, as well as, the initial distance of species traits from the community attractor. From relative abundance distributions of communities with different *d* (Fig 5a) (18), we see that the evenness of the distribution increases with *d*. We can directly relate this plasticity-driven increase in evenness to the amplitude and duration of fitness inequalities during transients (Fig 5b). As expected, the long-term community-level equilibrium leads to equal fitness among species. However, the magnitude and duration of transient fitness inequalities decrease with *d*. Fitness variance, which we define as variance in instantaneous growth rate accross species *σ*, increases initially due to the abundance of the total community increasing. However, the fitness variance begins to decrease once the community nears the supply vector and ecological equivalence is reached. The variance in equilibrium relative abundances in (Fig 5a) is a direct result of species traits during the transient, emphasizing the important role trait history plays in determining community structure. The transient time is not only controlled by *d* but also by how far away the community centroid initially is from the supply vector. We define a species sorting index, *SSI*, that captures the magnitude of trait-driven variation of relative abundances from an initially even species abundance distribution (equation (5)). We then calculate the initial centroid-supply vector distance, ∥*X*∥_2_, that captures how far in trait space a community begins from the attractor of the system (Fig 5c (red dashed line), equation (6)). We see that as *d* increases, the *SSI* is low (weak sorting) and independent from the initial centroid-supply vector distance (Fig 5d). As *d* decreases, we see that the *SSI* increases with ∥*X*∥_2_. We note that for very small ∥*X*∥_2_ no trait-driven variation in abundances is generated (Fig 5d). Transient time before ecological equivalence determines the trait-structure relationship, and transient time is given by initial centroid-supply distance divided by the speed a community acclimates, set by the plasticity rate. Hence, resource-use plasticity serves as a mechanism that can control the strength of the causal relationship between traits and community structure, as well as the slope of the rank abundance distribution.

**Fig. 5.**
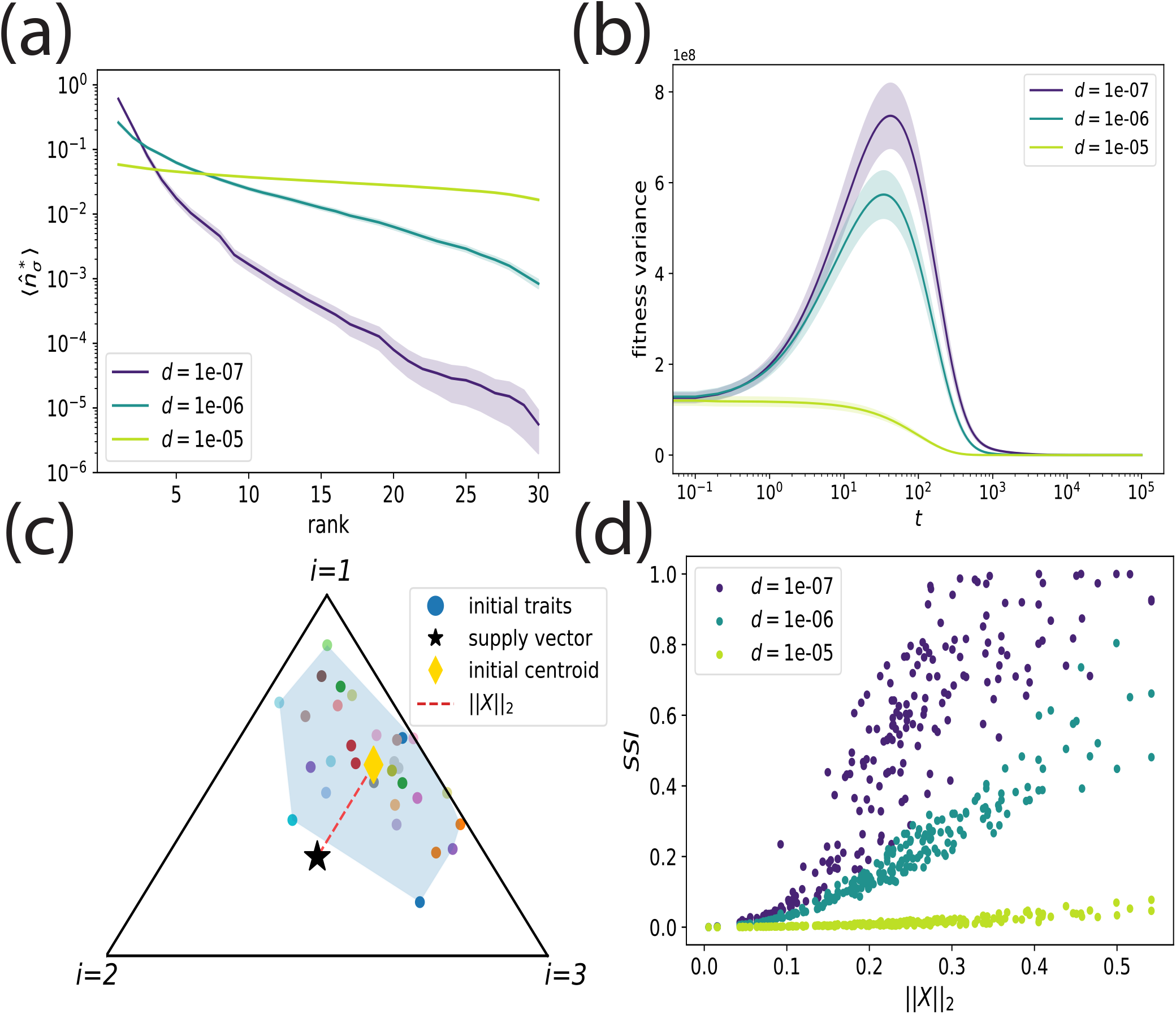
Plasticity rate *d* governs time-scale and magnitude of transient fitness inequalities that generate variation in species relative abundances. (a) Mean rank abundance distributions (with 95% confidence intervals) for different plasticity rates, *d*. (b) Time-series of fitness variance, i.e. variance in instantaneous growth rate, among species for different *d*. 95% confidence intervals are shown (c) Illustration of initial centroid-supply vector distance, ∥*X*∥_2_ (red dashed line). (d) Initial distance of community centroid to supply vector versus strength of ecological selection (trait-driven variation of relative abundances). Panel a, b and d result from 100 simulations for each plasticity rate with varying initial traits, *N*_*S*_ = 30, and *N*_*R*_ = 3. See SI for parameter details.

## Discussion

Community assembly and niche theory are typically understood as the result of sorting on heterogeneous and static traits among species, implying that trait changes occur over much slower timescales than ecological processes (15, 30, 31). However, trait and abundance dynamics can occur over similar timescales through selection (5, 32) and phenotypic plasticity (7, 23, 33, 34). Using a consumer-resource model, we show that plasticity in resource-use leads to a scale-dependent feedback between the dynamics of abundances and traits, with a complex, yet predictable relationship between species traits and their relative abundances. We first show how a single plastic species, while optimizing its own growth, can facilitate the coexistence of multiple competing species. The balance between selection on traits for minimizing interspecific competition and maximizing resource-use efficiency reveals that initial traits predict equilibrium species abundances better than equilibrium traits, uncovering equilibrium community structure’s dependence on transient traits. We further show that the temporal scale of resource use plasticity controls the strength of the causal relationship between traits and structure, and the maintenance of structure itself, by determining interspecific variation in abundance. We propose plasticity as a mechanism that determines the extent to which traits drive community structure by controlling the duration of transient dynamics before ecological equivalence is reached at equilibrium. The transient trait-structure coupling we report thus determines the strength of species sorting while predicting the equilibrium emergence of ecological equivalence from non-neutral trait-driven ecological processes. These results provide a new level of integration between niche and neutral frameworks through eco-evolutionary dynamics, and a mechanism for the generation of variation in community structure driven by species trait history.

### Plasticity and the emergence of keystone species

In the resource-competition model studied here, we show that plasticity is defined by the optimization of resource-use along two selection axes. The first is away from competitors, due to competition for the supplied resources. The second is towards the supply vector, which would be the optimal strategy in the absence of competitors. Here underlies a trade-off between avoiding competition (niche differentiation) and having the best suited strategy for that environment. We show how this environment-competition trade-off lets a single plastic species drive the coexistence of non-plastic species and the assembly of a community that would otherwise be reduced to at most a few species due to competitive exclusion. This is an interesting case of facilitation emerging from purely competitive interactions because the plastic species operates as a keystone species, due to it having a disproportionate effect on the rest of the community. The plastic species changes its consumption to avoid all the other competitors while getting closer to the supply vector. As a result, the force repelling the plastic strategy away from the competitors creates the flat fitness landscape at equilibrium that allows for coexistence through emergent ecological equivalence.

Our results highlight how competitive trade-offs that control selection on plastic traits can explain the emergence of facilitative interactions. Facilitative interactions among competitors have been shown to result from ecological processes such as spatial fluxes of organisms and resources (35, 36). They are also well studied as the result of evolutionary dynamics (6, 37). Rapid trait change in response to competitors has promoted coexistence in aquatic plant experiments where trait and abundance dynamics occurred on similar timescales (22, 23). In microbial communities, facilitation can occur via the cross-feeding of metabolic byproducts (31, 38, 39), which can be secreted due to a plastic phenomenon known as overflow metabolism (40). Incorporating trait plasticity to consumer-resource models with metabolic cross-feeding would bridge gaps between ecological, evolutionary, and intracellular processes.

In our study, the emergence of facilitation and of a keystone species can occur concurrently with competitive exclusion among non-plastic species and shape equilibrium community structure. During transient dynamics, the rescue effect of plasticity must be acting fast enough such that abundances of competitors are greater than zero before the community-level attractor is reached. Our results show how plasticity can result in the emergence of keystone species whose effect on community structure is predicted by their plasticity rate. Because coexistence can be driven by the plastic response of a single species, our results further provide a mechanism for the emergence of a keystone species that changes their traits to equalize fitness difference among species through their effect on resource availability.

#### Transient dynamics of traits and abundances can drive community structure

A core feature of this model is the strong relationship between equilibrium community structure and initial rather than final traits, which has been observed in experimental microbial systems (41, 42). In our work, this ’trait history’ phenomenon results from emergence of a ’community-level attractor’. At equilibrium, the abundance-weighted centroid of species’ traits converges to the supply vector. This centroid can be made up of infinite combinations of species’ traits and abundances meaning that different initial traits and abundances will not converge to a single fixed point. Additionally, we show that the transient dynamics that determine this equilibrium state are mediated by the rate of plasticity. When plasticity occurs quickly, the community equilibrium is reached much faster than species abundances, meaning that abundances are under weak sorting from traits for much of their dynamics. As a result, equilibrium community relative abundances are more related to initial abundances than to traits. This can be understood as a priority effect where early colonizers that reach high relative abundance early on (when resources are not limiting) retain their high relative abundance due to weak sorting. The consequence of this weak causal relationship between traits and abundance is that the species’ relative abundances depend on their initial values rather than being driven by trait-driven sorting. This is due to faster plasticity rates shortening the length of transients. When plasticity rate is slower, there is more variance in fitness that occurs over longer transients which leads to stronger species sorting. This stronger coupling between trait and abundance dynamics leads to uneven relative abundance distributions where species’ initial abundances no longer determine their equilibrium abundances.

Our results, which generate species-level variation depending on initial conditions, may explain the observed sensitivity of species-level composition to chance events and historical contingency observed in microbial communities(43–45). These studies observe a ’functional attractor’ at the family taxonomic-level while at the species level biological replicates show differences in equilibrium community structure, similar to the community-level attractor in our system. A recent experimental microcosm study links this functional attractor to resource use traits between families. One family metabolizes glucose through respiration and fermentation while the other family specializes in consuming organic acids solely through fermentation (43). The results from our study suggest that resource use plasticity may also explain community-level attractors with species-level variability of traits and abundances, implicating the role of transient dy-namics in determining community structure. Transient dynamics have become central to explaining major ecological (46), and eco-evolutionary phenomena (47). By incorporating phenotypic plasticity of resource consumption inspired by microbial systems, our work suggests a mechanism emerging from the coupling and eco-evolutionary transient dynamics of traits and abundances predicting when traits and abundances of initial colonizers play an important role in determining community structure.

### Revisiting the niche-neutral continuum with plasticity as a driver of community structure

Our work disentangles the dynamics of traits, abundances, and of their feedback as drivers of equilibrium community structure in competitive communities. We use an adaptive consumer-resource model where species, through plasticity, change their traits based on what combination of substitutable resources maximizes their growth. By tuning the plasticity rate, we can control the time scales of trait and abundance transient dynamics, during which fitness differences are generated by trait-resource suitability and lead to the production of interspecific variation in abundance. The transient time scale, which depends on plasticity rate and the initial distance of the community centroid to the supply vector, determines the extent to which traits drive community structure. At equilibrium, the community becomes fitness invariant due to the species driving resource concentrations to equal values, thus trait differences no longer generate fitness differences, and ecological equivalence emerges.

Similar models have been associated with neutral theory due to the similarity of their predictions of relative abundance distributions and to the emergence of fitness equivalence among species at equilibrium (15–17). However, these similarities are due to fundamentally different processes. In neutral theory, the steepness of relative abundance distributions is largely controlled by the rate at which new species are introduced to a local community either by immigration or speciation (4). In contrast, niche-based models typically predict equilibrium interspecific variation in relative abundance from fitness differences among species that are predictable from traits (3). In our system, fitness differences vanish at the community-level equilibrium, at which point, traits have no effect on structure. Instead, transient dynamics, whose timescales are largely governed by plasticity rate, are where trait-driven fitness differences generate and shape variation in community structure leading to relative abundance distribution patterns observed in nature. In other words, our results reveal mechanisms of neutral coexistence (traits do not affect abundance) and of variation in community structure (traits affect abundances) that are decoupled and partitioned between equilibrium and transient dynamics. Our study integrates niche-based and neutral explanations of community assembly via the trait-structure relationship, showing that predictions from both theories are compatible with niche-based processes that can drive community structure via a historical effect of transients and lead to equilibrium ecological equivalence. This reconciliation between niche-based community structure and ecological equivalence further provides a null model of community structure, predicting conditions for the emergence of interspecific variation in abundance: communities with an initially homogeneous distribution of abundance that exhibit fast plasticity and/or an initial community centroid near the supply vector, predict maximum equilibrium diversity with an even species abundance distribution (no community structure). Our results predict the emergence of community structure from (*i*) heterogeneous initial abundances that can for example be associated with colonization ability, and (*ii*) from trait dynamics occurring slower than abundance dynamics, or alternatively (*iii*) from communities not starting out pre-acclimated to their resource environments.

Our results illustrate the challenge of linking macro-ecological patterns to processes of community assembly (48, 49). We show how plasticity can drive the correspondence between equilibrium ecological equivalence and niche-based trait-abundance relationships in the absence of stochastic ecological drift. Our study proposes a theory of community structure and biodiversity driven by transient dynamics and equilibrium fitness equivalence, where relative abundance distributions are driven by transient eco-evolutionary trait-based processes. Further research on eco-evolutionary dynamics will likely illuminate other important roles of transient dynamics for community assembly.

