## Supplemental Information for "Resource-use plasticity governs the causal relationship between traits and community structure in model microbial communities"

### 1. SUPPLEMENTARY TEXT

#### A. Details of niche-based plasticity model

Here we show the derivation of the equation for trait dynamics. This growth rate optimization in equation 2 in the main text is implemented by gradient ascent. The time derivative of the resource uptake rates, $\dot{\alpha}_{\sigma i}$, follows the gradient ascent equation:

$$\dot{\alpha}_{\sigma i} = d\delta_\sigma \frac{\partial g_\sigma}{\partial \alpha_{\sigma i}} = d\delta_\sigma v_i r_i \tag{S1}$$

where $g_\sigma = \sum_i^{N_R} v_i \alpha_{\sigma i} r_i - \delta_\sigma$ and $r_i = r_i(c_i(t))$ for short. However, $\dot{\alpha}_{\sigma i}$, remains unbounded by the constraint $\sum_{i=1}^{N_R} \alpha_{\sigma i} = E_\sigma \; \forall \sigma$. Note that as opposed to in [1] we are using a strict constraint on $\alpha_{\sigma i}$ as opposed to an upper bound, assuming that all species are on their Pareto Frontier. We follow the Supplementary Information of [1] to derive our final equations with a few simplifications due to our strict constraint.

To ensure that the total resource uptake stays on the Pareto Frontier, the total uptake rate, $E_\sigma$, is constrained by the function:

$$\phi(\vec{\alpha}_\sigma) = \sum_{i=1}^{p} \alpha_{\sigma i} - E_\sigma = 0 \tag{S2}$$

We can project $\dot{\alpha}_{\sigma i}$ onto the tangent plane of $\phi(\vec{\alpha}_\sigma)$, ensuring the constraint is maintained, with the equation

$$\dot{\alpha}_{\sigma i} = d\delta_\sigma \frac{\partial g_\sigma}{\partial \alpha_{\sigma i}} - \frac{\nabla \phi(\vec{\alpha}_\sigma)}{\|\nabla \phi(\vec{\alpha}_\sigma)\|_2} \left( \frac{\nabla \phi(\vec{\alpha}_\sigma)}{\|\nabla \phi(\vec{\alpha}_\sigma)\|_2} \cdot d\delta_\sigma \frac{\partial g_\sigma}{\partial \alpha_{\sigma i}} \right) \tag{S3}$$

Our equation then becomes

$$\dot{\alpha}_{\sigma i} = d\delta_\sigma \frac{\partial g_\sigma}{\partial \alpha_{\sigma i}} - \frac{\frac{\partial \phi(\vec{\alpha}_\sigma)}{\partial \alpha_{\sigma i}}}{\sum_{\tau=1}^{N_S} \sum_{k=1}^{N_R} \left( \frac{\partial \phi(\vec{\alpha}_\sigma)}{\partial \alpha_{\tau k}} \right)^2} \left( \sum_{\rho=1}^{N_S} \sum_{j=1}^{N_R} \frac{\partial \phi(\vec{\alpha}_\sigma)}{\partial \alpha_{\rho j}} d\delta_\rho \frac{\partial g_\sigma}{\partial \alpha_{\rho j}} \right) \tag{S4}$$

This equation does not guarantee that the resource uptake rates remain non-negative. We modify so that $\alpha_{\sigma i}(t) \geq 0 \; \forall t$. This is done by introducing some auxiliary variables $\eta_{\sigma i}$ that the resource uptake rates are non-negative functions of, i.e $\alpha_{\sigma i} = F(\eta_{\sigma i})$ with $F(x) \geq 0 \, \forall x$. The dynamics for $\eta_{\sigma i}$ now become

$$\dot{\eta}_{\sigma i} = d\delta_\sigma \frac{\partial g_\sigma}{\partial \eta_{\sigma i}} - \frac{\partial \phi(\vec{\eta}_\sigma)/\partial \eta_{\sigma i}}{\sum_\tau^{N_S} \sum_k^{N_R} (\partial \phi(\vec{\eta}_\sigma)/\partial \eta_{\tau k})^2} \sum_\rho^{N_S} \sum_j^{N_R} \frac{\partial \phi(\vec{\eta}_\sigma)}{\partial \eta_{\rho j}} d\delta_\rho \frac{\partial g_\sigma}{\partial \eta_{\rho j}} \tag{S5}$$

If we change the variables and write this equation in terms of $\alpha_{\sigma i}$ we get

$$\dot{\alpha}_{\sigma i} = F'(\eta_{\sigma i})^2 \left[ d\delta_\sigma \frac{\partial g_\sigma}{\partial \alpha_{\sigma i}} - \frac{\partial \phi(\vec{\alpha}_\sigma)/\partial \alpha_{\sigma i}}{\sum_\tau^{N_S} \sum_k^{N_R} (F'(\eta_{\tau k}) \partial \phi(\vec{\alpha}_\sigma)/\partial \alpha_{\tau k})^2} \sum_\rho^{N_S} \sum_j^{N_R} \frac{\partial \phi(\vec{\alpha}_\sigma)}{\partial \alpha_{\rho j}} d\delta_\rho \frac{\partial g_\sigma}{\partial \alpha_{\rho j}} F'(\eta_{\rho j}) \right] \quad \text{(S6)}$$

Choosing $F(x) = x^2/4$ due always being positive and not generating any additional factors in the final equation, the dynamics become

$$\dot{\alpha}_{\sigma i} = \alpha_{\sigma i} \left[ d\delta_\sigma \frac{\partial g_\sigma}{\partial \alpha_{\sigma i}} - \frac{\partial \phi(\vec{\alpha}_\sigma)/\partial \alpha_{\sigma i}}{\sum_\tau^{N_S} \sum_k^{N_R} (\partial \phi(\vec{\alpha}_\sigma)/\partial \alpha_{\tau k})^2 \alpha_{\tau k}} \sum_\rho^{N_S} \sum_j^{N_R} \frac{\partial \phi(\vec{\alpha}_\sigma)}{\partial \alpha_{\rho j}} d\delta_\rho \alpha_{\rho j} \frac{\partial g_\sigma}{\partial \alpha_{\rho j}} \right] \quad \text{(S7)}$$

Finally, using $g_\sigma = \sum_i v_i \alpha_{\sigma i} r_i - \delta_\sigma$ and $\phi(\vec{\alpha}_\sigma) = \sum_i \alpha_{\sigma i}/E^*_\sigma - 1$, and solving the partial derivatives, the equation simplifies to

$$\dot{\alpha}_{\sigma i} = \alpha_{\sigma i} d\delta_\sigma \left[ v_i r_i - \frac{\sum_j v_j r_j \alpha_{\sigma j}}{\sum_k^{N_R} \alpha_{\tau k}} \right] \quad \text{(S8)}$$

Thus, the correct form has $\alpha_{\sigma k}$ in the denominator of the last term, ensuring the units of the two two terms balance correctly. We then to simplify to

$$\dot{\alpha}_{\sigma i} = \alpha_{\sigma i} d\delta_\sigma \left( v_i r_i - \frac{\sum_{j=1}^{N_R} v_j r_j \alpha_{\sigma j}}{E_\sigma} \right) \quad \text{(S9)}$$

where $r_i = r_i(c_i(t))$ for short. Our final system of equations becomes:

$$\begin{aligned} \frac{dn_\sigma}{dt} &= n_\sigma \left( \sum_i^{N_R} v_i \alpha_{\sigma i} r_i - \delta_\sigma \right) \\ \frac{dc_i}{dt} &= s_i - r_i \sum_\sigma^{N_S} \alpha_{\sigma i} n_\sigma \\ \frac{d\alpha_{\sigma i}}{dt} &= \alpha_{\sigma i} d\delta_\sigma \left( v_i r_i - \frac{\sum_{j=1}^{N_R} v_j r_j \alpha_{\sigma j}}{E_\sigma} \right) \end{aligned} \quad \text{(S10)}$$

### B. Details on statistical model

Our statistical model (equation (4) in main text), assumes equilibrium relative abundances can be determined by traits due to growth maximization by plasticity. Plastic species attempt to maximize trait distance from competitors and minimize trait distance to supply vector. We show a more general version of equation 4 in the main text where these distances are taken as functions of time

$$\log(n^*_\sigma) = a \frac{\sum_{k=1}^{N_S} A_{\sigma k}(t)}{N_S - 1} - b\vec{\phi}_\sigma(t) \quad \text{(S11)}$$

where parameters $a$ and $b$ are fitted by ordinary least squares regression. We first show the values of the coefficients $a$ and $b$ which we refer to as effect size (Fig (S2a)) as a function of plasticity rate, $d$. We see that supply distance is more important in determining equilibrium relative abundances than competitor distance and that the relative importance of these distances does not depend on plasticity rate. Here we take effect sizes from our statistical model with input distances calculated at $t = 0$. Additionally, if we predict equilibrium relative abundance distribution from $A_{\sigma k}(t)$ and $\vec{\phi}_\sigma(t)$ throughout a time-series, we see that the sharp decrease in the variance explained from the total model comes from supply distance $\vec{\phi}_\sigma(t)$ (Fig (S2a)). From this we can see that as the weighted community centroid gets closer to the attractor, $\hat{s}_i$, less variance is explained from traits. We use $R^2$ as the performance metric for variance explained. Here $N_S = 10$, $N_R = 3$, and $d = 5 \times 10^{-7}$. The rest of the parameters being the same or drawn from the same distributions as the Figure 3 in the main text.

#### C. Ecological equivalence emerging from consumer-resource model with trade-offs

Ecological equivalence emerges in this metabolic trade-off model [2] due to ratio of the total metabolic rate, $E_\sigma$ and the maintenance rates, $\delta_\sigma$, being equal to the same constant $Q = \frac{E_\sigma}{\delta_\sigma}\ \forall\sigma$. The metabolic theory of ecology argues that both $E_\sigma$ and $\delta_\sigma$ are proportional to the characteristic mass, $M_\sigma$, of species $\sigma$ to the power of a universal exponent, $\lambda$ [3, 4]. Let's say we introduce biomass, $B_\sigma(t)$, as a state variable related to abundance by $B_\sigma(t) = M_\sigma n_\sigma(t)$. Then by the metabolic theory of ecology $E_\sigma \propto M_\sigma^\lambda$ and $\delta_\sigma \propto M_\sigma^\lambda$, thus

$$\frac{E_\sigma}{\delta_\sigma} = \frac{M_\sigma^\lambda}{M_\sigma^\lambda} = Q \tag{S12}$$

[1]. These rates also depend on temperature, however, we assume temperature to be constant in a local environment of competing species. This is analogous to a perfect trade-off between fecundity and mortality where neutral coexistence is maintained by species driving resource levels to equal values at equilibrium (i.e. having equivalent $R^*$'s [5]).

#### D. Magnitude of transient fitness variance depends on $N_R$

The magnitude and duration of transient fitness differences depends on if the normalized supply vector, $\hat{s}_i$, begins in the initial convex hull of the normalized resource uptake rates, $\hat{\alpha}_{\sigma i}(t=0)$. The probability that this occurs of course depends on the distributions that $\hat{s}_i$ and $\hat{\alpha}_{\sigma i}(t=0)$ are drawn from but greatly depends on the number of resources, $N_R$. Let's assume that both $\hat{s}_i$ and $\hat{\alpha}_{\sigma i}(t=0)$ are drawn from uniform distributions such that $\sum_{i=1}^{N_R} \hat{\alpha}_{\sigma i} = 1$ and $\sum_{i=1}^{N_R} \hat{s}_i = 1$. This is a special case of the Dirichlet distribution where its $N_R$ parameters are all 1, i.e. $Dir(1, ..., 1_{N_R})$. Thus the probability that $\hat{s}_i$ is in the convex hull of $\hat{\alpha}_{\sigma i}(t=0)$ is given by the $N_R - 1$ dimensional volume of the convex hull divided by the $N_R - 1$ dimensional volume of the entire simplex. We sample 1000 convex hulls made from $\hat{\alpha}_{\sigma i}(t=0)$ for different combination of $N_R$ and $N_S$ and show the average probabilities of $\hat{s}_i$ being in the trait convex hull (Fig. S3). We see that these probabilities increase with $N_S$ while sharply decrease with $N_R$ (Fig. S3a). We show an example of communities with $N_S = 30$ and $N_S = 3$ to demonstrate that the average sorting ($SSI$) is stronger when the convex hull condition is not originally met (Fig. S3b). We calculate convex hull volumes with the `ConvexHull` function in the package `scipy.spatial`.

### 2. SIMULATION DETAILS

Many of the parameters in niche-based plasticity model were kept to be the same across species. We mostly changed plasticity rate, $d$, the initial resource-use rates $\alpha_{\sigma i}(t=0)$, and the supply rates $s_i$. We drew initial traits from the Dirichlet distribution $Dir(\mathcal{U}\{1,5\})$. Additionally, in the main text we often show abundances, traits, and supply rates in relative terms which are unit-less. However, these terms keep the same units as in [1]. Although $d$ is not technically a rate, $d\delta_\sigma$, is. Since $\delta_\sigma$ is the same across species in our simulations, we use $d$ to control its scale. Thus it acts as a control on the rate species can change their resource-use strategies. For simplicity and intuition's sake we refer to $d$ as plasticity rate in the text.

##### Figure 2

$v_i = 1\times10^8 \frac{\text{cells}}{\text{g of resource}} \forall i$, $\delta_\sigma = 5\times10^{-3} \frac{1}{\text{hour}} \forall\sigma$, $Q = 1\times10^{-4} \frac{\text{g of resource}}{\text{cell}}$, $E_\sigma = 5\times10^{-7} \frac{\text{g of resource}}{\text{cells}\cdot\text{hour}} \forall\sigma$, $K_i = 1\times10^{-5} \frac{\text{g of resource}}{\text{mL}} \forall i$, $s_i \sim \mathcal{U}(1\times10^{-6}, 1\times10^{-2}) \frac{\text{g of resource}}{\text{mL}\cdot\text{hour}}$, $n_\sigma(t=0) = 1\times10^6 \frac{\text{cells}}{\text{mL}} \forall\sigma$, $c_i(t=0) = 1\times10^{-3} \frac{\text{g of resource}}{\text{mL}} \forall i$, $\alpha_{\sigma i}(t=0) \sim Dir(\mathcal{U}\{1,5\}) \frac{\text{g of resource}}{\text{cell}\cdot\text{hour}}$. In 2b $d = 0 \frac{\text{g of resource}}{\text{cell}} \forall\sigma$ while in 2c $d = 6\times10^{-7} \frac{\text{g of resource}}{\text{cell}}$ for $\sigma = 1$ and 0 for all other species. $N_S = 10$ and $N_R = 3$.

##### Figure 3

$v_i = 1\times10^8 \frac{\text{cells}}{\text{g of resource}} \forall i$, $\delta_\sigma = 5\times10^{-3} \frac{1}{\text{hour}} \forall\sigma$, $Q = 1\times10^{-4} \frac{\text{g of resource}}{\text{cell}}$, $E_\sigma = 5\times10^{-7} \frac{\text{g of resource}}{\text{cells}\cdot\text{hour}} \forall\sigma$, $K_i = 1\times10^{-5} \frac{\text{g of resource}}{\text{mL}} \forall i$, $s_i \sim \mathcal{U}(1\times10^{-6}, 1\times10^{-2}) \frac{\text{g of resource}}{\text{mL}\cdot\text{hour}}$, $n_\sigma(t=0) = 1\times10^6 \frac{\text{cells}}{\text{mL}} \forall\sigma$, $c_i(t=0) = 1\times10^{-3} \frac{\text{g of resource}}{\text{mL}} \forall i$, $\alpha_{\sigma i}(t=0) \sim Dir(\mathcal{U}\{1,5\}) \frac{\text{g of resource}}{\text{cell}\cdot\text{hour}}$, and $d = 5\times10^{-7} \frac{\text{g of resource}}{\text{cell}}$. $N_S = 10$ and $N_R = 3$.

**Figure 4**

$v_i = 1 \times 10^8 \frac{\text{cells}}{\text{g of resource}} \forall i$, $\delta_\sigma = 5 \times 10^{-3} \frac{1}{\text{hour}} \forall \sigma$, $Q = 1 \times 10^{-4} \frac{\text{g of resource}}{\text{cell}}$, $E_\sigma = 5 \times 10^{-7} \frac{\text{g of resource}}{\text{cells}\cdot\text{ hour}} \forall \sigma$, $K_i = 1 \times 10^{-5} \frac{\text{g of resource}}{\text{mL}} \forall i$, $s_i \sim \mathcal{U}(1 \times 10^{-5}, 1 \times 10^{-2}) \frac{\text{g of resource}}{\text{mL}\cdot\text{hour}}$, $n_\sigma(t=0) \sim \mathcal{U}(1 \times 10^3, 1 \times 10^6) \frac{\text{cells}}{\text{mL}} \forall \sigma$, $c_i(t=0) = 1 \times 10^{-3} \frac{\text{g of resource}}{\text{mL}} \forall i$, $\alpha_{\sigma i}(t=0) \sim Dir(\mathcal{U}\{1,5\}) \frac{\text{g of resource}}{\text{cell}\cdot\text{ hour}}$. $N_S = 30$ and $N_R = 3$.

**Figure 5**

$v_i = 1 \times 10^8 \frac{\text{cells}}{\text{g of resource}} \forall i$, $\delta_\sigma = 5 \times 10^{-3} \frac{1}{\text{hour}} \forall \sigma$, $Q = 1 \times 10^{-4} \frac{\text{g of resource}}{\text{cell}}$, $E_\sigma = 5 \times 10^{-7} \frac{\text{g of resource}}{\text{cells}\cdot\text{ hour}} \forall \sigma$, $K_i = 1 \times 10^{-5} \frac{\text{g of resource}}{\text{mL}} \forall i$, $s_i \sim \mathcal{U}(1 \times 10^{-5}, 1 \times 10^{-2}) \frac{\text{g of resource}}{\text{mL}\cdot\text{hour}}$, $n_\sigma(t=0) = 1 \times 10^6 \frac{\text{cells}}{\text{mL}} \forall \sigma$, $c_i(t=0) = 1 \times 10^{-3} \frac{\text{g of resource}}{\text{mL}} \forall i$, $\alpha_{\sigma i}(t=0) \sim Dir(\mathcal{U}\{1,5\}) \frac{\text{g of resource}}{\text{cell}\cdot\text{ hour}}$. $N_S = 30$ and $N_R = 3$.

**Figure SI 1**

$N_S = 10$, $N_R = 3$, and $d = 5 \times 10^{-7}$. All other parameters are the same as in Figure 3 in the main text.

**Figure SI 2**

$N_S = 10$, $N_R = 3$, and $d = 5 \times 10^{-7}$. All other parameters are the same as in Figure 3 in the main text.

**Figure SI 3**

All other parameters not explicitly changed in the figure are the same as in Figure 3 in the main text. In Figure (S3b) $N_S = 30$, and $N_R = 3$

### 3. FIGURES

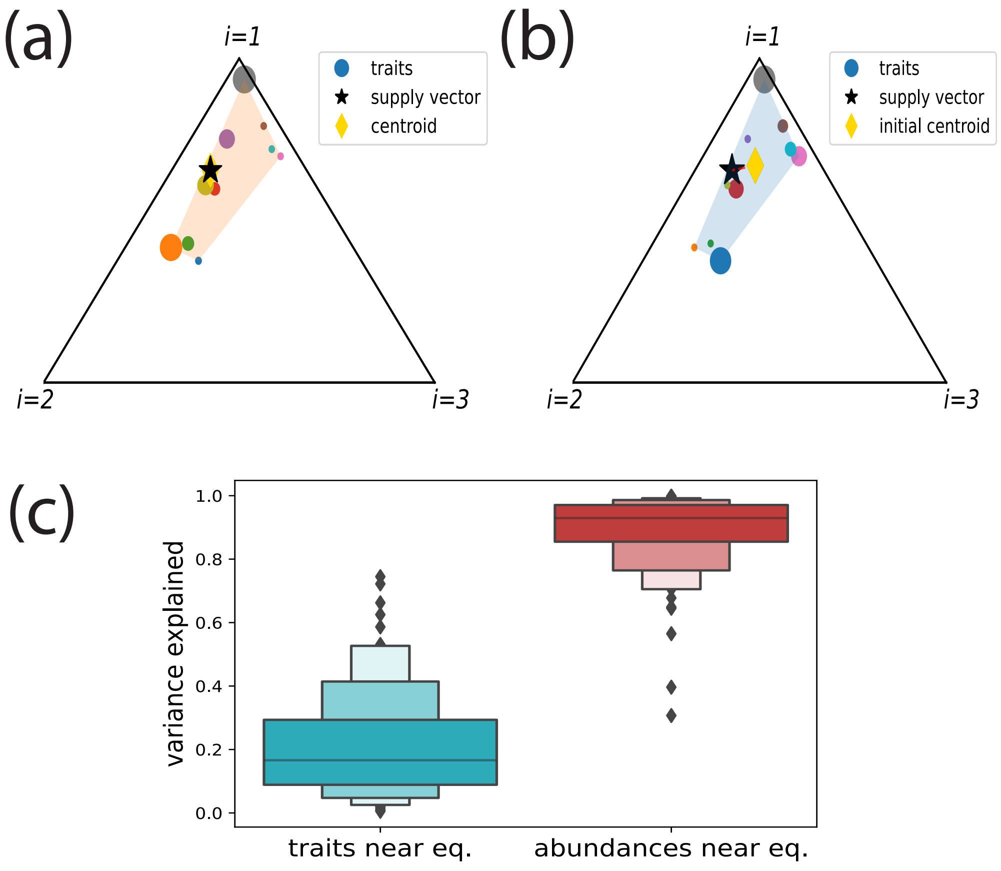


**Fig. S1.** Abundance shuffling experiment. (a) Simplex diagram of at equilibrium (orange) showing equilibrium traits $\hat{\alpha}^*_{\sigma i}$ (dots), supply vector $\hat{s}_i$ (black star), and centroid $\|X\|_2$ (yellow diamond). (b) Simplex diagram after perturbation by randomly shuffling abundances onto new traits and resulting new (initial) weighted-centroid. (c) the variance explained ($R^2$) of traits and abundances after shuffling perturbation.

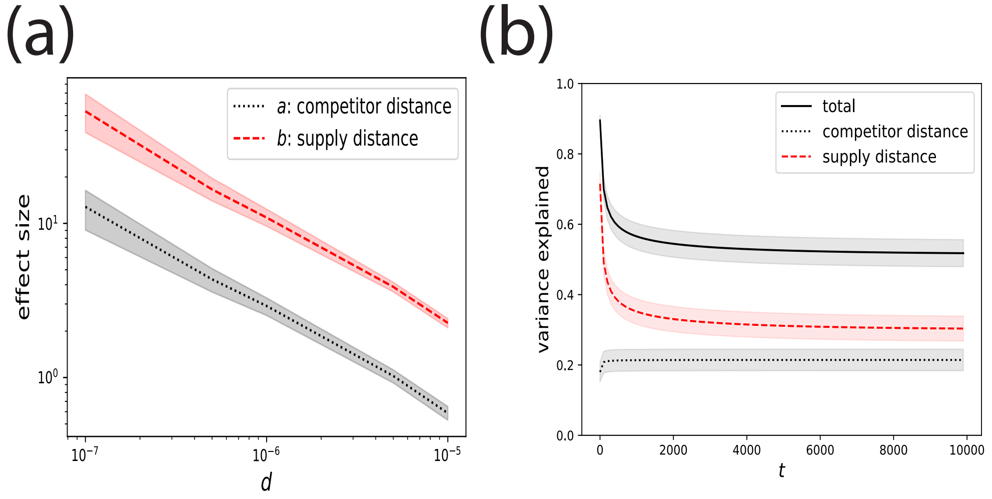


**Fig. S2.** Breakdown of biotic and abiotic terms in statistical model (a) effect size $a$ of competitor distance (black dotted), and effect size $b$ of supply distance (red dashed) (from equation 3 in main text). Distances are calculated from initial traits $\alpha_{\sigma i}(t = 0)$ (b) variance explained in equilibrium community structure as a function of traits through for total model (black solid line), only competitor distance (black dotted), and only supply distance (red dashed).

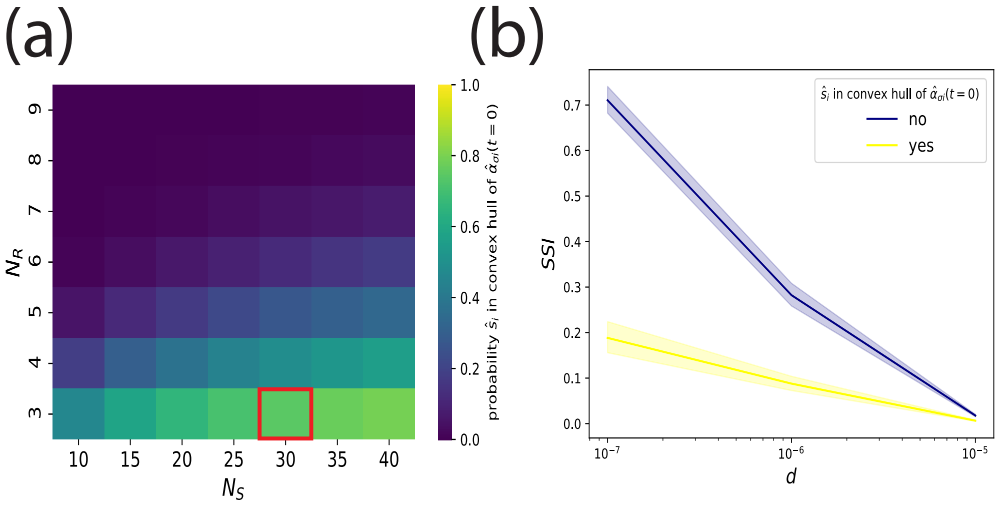


**Fig. S3.** Probability and outcome of supply vector, $\hat{s}_i$, being in convex hull of initial traits, $\hat{\alpha}_{\sigma i}$. (a) Probability of $\hat{s}_i$ being in convex hull of $\hat{\alpha}_{\sigma i}$ for different $N_R$ and $N_S$. (b) Communities of $N_S = 30$ and $N_R = 3$ (outlined in red from (a)) showing species sorting index, $SSI$, as a function of plasticity rate $d$ when $\hat{s}_i$ is or is not in convex hull of $\hat{\alpha}_{\sigma i}$. 95% confidence interval shown.
